# Tracking interannual temperature variation: phenological plasticity and reproductive stability in tropical hornbills

**DOI:** 10.64898/2026.05.27.728085

**Authors:** Aparajita Datta, Ritobroto Chanda, Rohit Naniwadekar, Karishma Pradhan, Soumya Banerjee, Khem Thapa, Jitendra Brah

## Abstract

Climate change in the tropics is expected to manifest differently than in temperate regions: rather than directional warming, its ecological signature may be dominated by increased year-to-year variability against a backdrop of comparatively modest average change. This distinction matters for organisms that evolved in historically stable climates and may respond strongly to year-to-year fluctuations even in the absence of long-term trends. Whether tropical birds track this kind of interannual variability, and whether doing so has an influence on nesting success, remains poorly understood, particularly for cavity-nesting frugivores whose breeding is tightly bound to seasonal resource availability. We used 27 years of nest monitoring data on three sympatric hornbill species at Pakke Tiger Reserve, northeast India, to examine long-term changes in nest-entry timing, nest occupancy, nesting success, and breeding duration, and to test whether interannual variation in local climate predicted breeding parameters. Local temperature and rainfall varied substantially among years without clear long-term trends. Nest-entry timing remained stable in Great and Wreathed Hornbills but became later in Oriental Pied Hornbills; Wreathed Hornbill nest occupancy also declined, whereas nesting success showed no directional change. Across the three species, warmer pre-breeding conditions were associated with earlier nest entry, while rainfall added no independent explanatory power once temperature was accounted for. The three species did not differ detectably in their sensitivity to temperature, despite differences in body size and breeding-cycle length. Our results illustrate how tropical cavity-nesting frugivores may track interannual climate variation without immediate breeding costs, although the strength of support for the temperature association differs among species and climate change may manifest primarily as altered variability rather than strong directional trends.

## 1. Introduction

Predicting species’ responses to climate change requires understanding not only whether populations can track environmental shifts phenologically, but also whether such plasticity translates into sustained reproductive and demographic performance, a relationship that remains poorly resolved for tropical species (Cannone et al. 2022; Vega et al. 2021). Climate change and increased climate variability pose multifaceted threats to avian breeding systems, yet the specific mechanisms and manifestations of these impacts remain poorly understood, particularly in tropical regions (Levillain et al. 2025; Brawn et al. 2017). Understanding how climate variability affects vulnerable bird species requires examining not only shifts in breeding timing but also broader changes in breeding ecology (Descamps et al. 2014). While much attention has focused on shifts in the timing of breeding like earlier nest initiation or prolonged breeding seasons (Visser & Both 2005), emerging evidence suggests that climate impacts on breeding ecology extend well beyond phenological timing (Whitenack et al. 2023). Birds use a variety of nesting structures and sites. While some species build their own nests, others rely on existing structures, with secondary cavity-nesting birds being particularly constrained by the availability of suitable nest cavities. Changes in breeding parameters such as nesting occupancy rates (the proportion of available and suitable nesting cavities occupied), nesting success (fledging success), nesting duration, and overall chick production are likely to be equally sensitive to climate variability (Arct et al. 2025). Responses vary considerably: some populations show delayed breeding or increased temporal plasticity (Senapathi et al. 2011; Barve et al. 2025), others non-linear threshold responses (Andreasson et al. 2023), and recent work in Andean cloud forests shows drought-induced breeding skips in insectivorous birds (Newell et al. 2026). Extreme weather events can disrupt nest initiation independently of seasonal trends (Taff and Shipley 2023).

Climate change is not experienced uniformly across latitudes. In temperate systems, ecological responses to climate change are frequently framed around directional shifts — progressively earlier springs and steadily rising mean temperatures (Parmesan & Yohe 2003; Root et al. 2003) — because that is the dominant signal in these regions. Tropical climate change looks different: mean warming is often comparatively modest, but organisms there evolved against a backdrop of low historical climate variability, so even non-directional, erratic fluctuations can carry disproportionate biological consequences (Deutsch et al. 2008; Vasseur et al. 2014), with breeding failures in birds often concentrated in extreme years rather than tracking a steady linear trend (Ventura et al. 2023). Recent global analyses suggest that tropical ecosystems are likely to experience climatic conditions that are increasingly outside the range of their historical variability, with tropical regions among those projected to encounter novel climatic conditions particularly rapidly (Trisos et al. 2020). A directional-trend framework borrowed from temperate systems may therefore be poorly suited to understanding how tropical species respond to a changing climate, and the more relevant question may not be whether conditions are trending in one direction, but whether organisms track the year-to-year variability itself, and whether doing so carries a breeding cost.

Most empirical evidence for climate-driven phenological shifts comes from temperate systems, where migratory insectivorous birds like pied flycatchers and great tits show breeding mismatches between breeding timing and prey availability due to differential warming responses (Visser et al. 1998; Both & Visser 2001; Dunn 2004; Källander et al. 2017). When birds’ phenological shifts lag behind their food resources, nesting success and population viability decline (Visser & Both 2005; Glądalski et al. 2022). However, this knowledge base is shaped by both strong geographic and taxonomic biases and a focus on a few particular response variables. Most studies focus on temperate insectivores, where long-term datasets document earlier egg-laying in response to warming (Dunn and Møller 2019; Halupka et al. 2023), but fewer examine broader breeding parameters such as nest occupancy, success, or breeding duration (Telenský et al. 2020). Studies specifically examining climate-phenology relationships in frugivorous birds remain limited across both temperate and tropical systems, and existing work suggests that food availability and climate can jointly influence the timing of avian activity (Kanerva et al. 2020; Newell et al. 2026). Phenological responses to climate variation in tropical frugivorous and cavity-nesting birds remain poorly documented, particularly over long time periods, and the links between climatic conditions, food resources, and breeding remain unclear. Existing studies span temperate and tropical systems and show that breeding timing and reproductive performance can respond to climatic variation, but responses are variable among species and systems (Drake & Martin 2018; Liu et al. 2021; Rush et al. 2024). This leaves limited understanding of how tropical frugivores respond to interannual climate variation and whether changes in breeding timing translate into changes in other components of reproductive performance.

This gap is particularly critical because tropical species may exhibit fundamentally different climate sensitivities than temperate counterparts, reflecting distinct thermal physiology, strong dependence on pulsed local resources (fruiting regimes), and varying phenological capacities across monsoon- vs equatorial systems (Pollock et al. 2020; Reich 1995). Recent global syntheses from the tropics further suggest that classical temperate paradigms may not hold, and that demographic responses can differ systematically between temperate and tropical systems (Smart et al. 2024).

Hornbills (Bucerotidae) are large, long-lived secondary cavity-nesting birds occurring in tropical Asia and Africa. Asian forest hornbills are generally highly frugivorous, and their breeding phenology is intimately linked to fruit availability and climate (Datta & Rawat 2003; Kinnaird & O’Brien 2007). Their breeding strategy, in which females are sealed in cavities and depend entirely on male provisioning throughout a prolonged incubation and chick-rearing period, makes them vulnerable to resource scarcity during fruit shortages and to thermal stress from climate change. Climate warming and increasing extremes are widely linked to reduced avian reproductive performance (Cunningham et al. 2013; Conradie et al. 2019).

In African hornbills, high temperatures and low rainfall have been shown to reduce nesting success and alter breeding onset (van de Ven et al. 2021; Pattinson et al. 2022), although these studies largely relied on artificial nest boxes, which can differ substantially from natural tree cavities in their thermal environments, with natural cavities providing greater buffering against ambient temperature extremes (Pattinson et al. 2022; Combrink et al. 2017). Observations from East African hornbills also suggest that prolonged drought can be associated with unusual out-of-season breeding attempts, indicating that climatic extremes may affect breeding timing as well as success (O’Brien & Kinnaird 2022). In Asia, climate change is also expected to reshape hornbill distributions and breeding conditions across regions with very different climatic regimes, from strong monsoon seasonality to more aseasonal tropical environments (Numata et al. 2022; Bajagain et al. 2025). However, despite growing work on hornbill ecology and climate-related changes in breeding, direct evidence on how climate interacts with nest occupancy, timing, and success in Asian forest hornbills remains scarce.

Here we utilize data from a 27-year (1998-2025) nest monitoring project for three sympatric hornbill species (Great Hornbill *Buceros bicornis*, Wreathed Hornbill *Rhyticeros undulatus*, Oriental Pied Hornbill *Anthracoceros albirostris*) from a Protected Area in the Eastern Himalaya to examine how climate variability shapes breeding ecology in threatened tropical cavity-nesting frugivores. This long-term dataset provides a rare opportunity in a tropical forest system to distinguish directional changes over decades from responses to interannual climate variability, and to examine whether phenological plasticity is associated with stability or variation in reproductive performance.

Specifically, we test the following questions:

1. Have hornbill breeding parameters (nest entry timing, nest occupancy, nesting success, breeding duration) changed over the past 27 years (1998-2025), and are observed patterns driven by long-term climate trends or interannual climate variability? We hypothesize that breeding parameters will vary primarily with interannual climate variability rather than follow progressive directional trends, with year-to-year variation potentially driven by local climate fluctuations, including ENSO.
2. How do temperature and rainfall during different phases of the breeding cycle (pre-breeding, early breeding, peak breeding) and over different pre-entry windows explain interannual variation in nest entry timing, nest occupancy, nesting success and breeding duration? We hypothesize that warmer pre-breeding conditions will be associated with earlier nest entry. Given differences in body size and breeding-cycle duration among the three species, we also test whether the strength of this association differs among species.

## 2. Methods

### 2.1 Study Site

We conducted this study at Pakke Tiger Reserve (862 km^2^, 92°36’–93°09’E, 26°54’–27°16’N) in western Arunachal Pradesh, northeast India, which is part of the Eastern Himalaya Biodiversity Hotspot (Mittermeier et al. 1999). The reserve ranges in elevation from 100 to 1,700 m above sea level. The dominant forest type is Assam Valley tropical semi-evergreen forest (Champion and Seth 1968), characterized by multi-storied canopy structure with major plant families including Lauraceae, Euphorbiaceae, Myrtaceae, and Meliaceae. Common tree species include *Monoon simiarum* (Annonaceae), *Pterospermum acerifolium* (Malvaceae), *Pterygota alata* (Malvaceae), and *Duabanga grandiflora* (Lythraceae), with emergent species including *Tetrameles nudiflora* (Tetramelaceae), *Ailanthus integrifolia* (Simaroubaceae), and *Liquidambar excelsa* (Altingiaceae) (Page et al. 2022).

The reserve is located at 27°N in the Eastern Himalaya monsoon zone and experienced modest warming (mean maximum temperature increase of 1.2°C between 1983-1995 and 2011-2023 baselines; Datta et al. 2025). Pakke Tiger Reserve was free from anthropogenic disturbances that could affect hornbill nesting during the entire study period (1998-2025), ensuring that observed patterns in hornbill nest entry timing, nest occupancy, nesting success, and breeding duration reflect natural environmental variation alone.

### 2.2 Study Species

We monitored a total of 65 nests of three sympatric hornbill species: Great Hornbill (*Buceros bicornis*), Wreathed Hornbill (*Rhyticeros undulatus*), and Oriental Pied Hornbill (*Anthracoceros albirostris*) that are long-lived secondary cavity-nesters with a highly specialized breeding strategy. Females seal themselves inside natural tree cavities during breeding, depending entirely on male provisioning throughout the incubation and chick-rearing period for 3-4 months. This extended period of cavity occupancy makes hornbills uniquely vulnerable to resource fluctuations and thermal stress. All three species are primarily frugivorous, with breeding phenology tightly linked to fruit availability in tropical forests (Datta 2001; Datta & Rawat 2003; Naniwadekar et al. 2015; Poonswad et al. 2004). The three species differ in body size, with Great Hornbill being the largest (2.6 to 3.9 kg, 120-150 cm), followed by Wreathed Hornbill (2-3.6 kg, 100-117 cm) and Oriental Pied Hornbill (680-907 gm, 70-85 cm) (Kemp 1995; Birds of the World, 2026), suggesting potential variation in sensitivity to climate. There is sexual dimorphism in body size with females being smaller (Kemp 1995).

The hornbill breeding season in this region is usually from March to August, while from January to February, hornbill pairs are seen actively searching for, visiting, inspecting, cleaning nest cavities (Datta 2001; Datta & Rawat 2004). Courtship and mating behaviours are also observed in the pre-breeding period between adult pairs, often at nest trees.

### 2.3 Nest Monitoring

Nest trees with cavities used by hornbills for breeding were identified through surveys of the study area conducted from 1997 onwards. Identified nest trees were tagged, numbered and locations were recorded. These nest trees were monitored as part of the long-term nest monitoring program regardless of nest occupancy status in any given year.

We monitored hornbill breeding activity from 1998 to 2025, with some modifications to protocol over this period. From 1998 to 2000, nest monitoring was conducted systematically as part of a detailed study on hornbill breeding biology (Datta 2001). Hornbill nest monitoring was re-initiated in 2003 and till 2006 was carried out opportunistically with varying intensity. In 2007, we implemented a standardized monitoring protocol maintained consistently through 2025 by 2 to 4 trained observers per season, with annual training refreshers and data quality checks ensuring methodological consistency across field personnel changes. Due to this change, species-specific nest occupancy and success analyses use 2007-2025 data, while overall trends span 1998-2025. Nest entry timing was analysed using the full available record (1998-2025). The study focused on threatened hornbill species (which are accorded the highest legal protection status) inside a protected tiger reserve, monitoring was carried out under permits issued by the Arunachal Pradesh Forest Department and was non-invasive and observational. All monitoring followed standardized nest observation protocols minimizing disturbance to occupied nests (Barve et al. 2020a, b).

During the pre-breeding and breeding season (February-August), trained observers monitored known hornbill nest cavities to record nest visits, nest entry dates, nest occupancy breeding outcomes, female and chick exit dates. We defined nest entry as the first date on which a female was observed inside the cavity. Females typically begin cavity entry before the entrance is fully sealed, and sealing may occur over several subsequent days. Observers recorded the progressive sealing of the entrance, including observations of partially or fully sealed entrances, but these later observations were not used to redefine the nest entry date. Nest occupancy was determined from sealed nest entrances, male provisioning at the cavity, or vocalizations from within the cavity. The exit date was the day the fledged chicks emerged from the cavity. Nests were typically checked every 2-3 days; when the exact entry or exit day could not be observed directly, we estimated the date as the midpoint between the last “pre-event” and first “post-event” check. Nests with monitoring gaps > 4 days around entry or exit times were excluded from entry-timing and nesting duration analyses. Nesting duration (length of the breeding cycle) was calculated only for nests with both usable entry and exit dates.

Nesting success was defined as fledging of at least one chick, based on direct observation of fledging or chicks seen or heard near the nest tree with attending parents. When a 2-4 day monitoring gap spanned a potential exit event, and if the nest had been active during previous monitoring visits till the end and the nest showed no evidence of failure, we classified it as successful. Published studies report generally low nest predation rates in hornbills, including at our study site, with occasional predation by Yellow-throated Martens (*Martes flavigula*) documented during our long-term monitoring (three instances; AD/KT pers. obs.; Lahiri 2017) and by Binturongs (*Arctictis binturong*) reported elsewhere in Thailand (Poonswad et al. 1999; Kinnaird & O’Brien 2007). Given the rarity of these observations, we consider any positive bias in our estimates of nesting success due to undetected predation to be small.

Nest monitoring gaps occurred in 2001–2002 (no data), 2004 (partial success data missing), 2011 (success data missing due to monitoring constraints), and February 2020-April 2021 (COVID-19 pandemic and permit restrictions).

We calculated the following annual breeding parameters:

- Nest occupancy: proportion of monitored usable cavities occupied per species per year.
- Nesting success: proportion of occupied nests fledging at least one chick.
- Nest entry timing: median day of year (DOY) of female cavity entry per species per year.
- Breeding duration: mean number of days from nest entry to exit per species-year

### 2.4 Climate Data Collection

Historical temperature and rainfall records from 1983-1995; 1997-2000 (Tipi Orchid Research Centre; Datta 2001) were compiled for longer-term context. .We established a weather station (H21-USB data logger) in Seijosa, at the edge of Pakke Tiger Reserve (∼1 km from the boundary), in April 2011. The station recorded temperature (°C) and rainfall (mm) hourly. Raw climate data were processed up to 31 December 2025, with temperature and rainfall data cleaned independently before analysis. We processed raw hourly climate data into daily and seasonal summaries for analysis. Daily values were calculated as mean temperature (mean of all hourly readings) and total daily rainfall (sum of hourly readings). Based on long-term observations of hornbill breeding for three sympatric species at Pakke, nest-entry dates generally span February to early May, with most nest entries occurring from late February through April. We therefore defined a pre-breeding period preceding the main nest-entry period, followed by early and peak breeding periods, and used fixed calendar windows to represent these stages:

- Pre-breeding period (DOY 20-59; January 20-February 28): Pre-breeding period when nest initiation decisions occur
- Early breeding period (DOY 60-120; March 1-April 30): Nest building, egg-laying, and early incubation
- Peak breeding period (DOY 121-181; May 1-June 30): Late incubation and chick-rearing

For each window, we calculated mean temperature and total rainfall per year. We additionally obtained the Multivariate ENSO Index version 2 (MEI v2) from the NOAA Physical Sciences Laboratory (Wolter & Timlin 2011; NOAA PSL). For each breeding year, pre-breeding MEI was calculated as the mean of November and December of the preceding year and January and February of the breeding year, provided that values were available for at least three of these four months.

### 2.5 Statistical Analyses

All analyses were conducted in R version 4.3.1 (R Core Team 2024) using RStudio 2025.09.1+401. We set the significance threshold at α = 0.05 for all statistical tests.

#### 2.5.1 Long-term Trends in Breeding Parameters

We examined temporal trends in hornbill breeding parameters across multiple time periods based on data availability and monitoring standardization. Overall nest occupancy and nesting success were analysed from 1998 to 2025. Species-specific analyses of nest occupancy and nesting success used data from 2007 to 2025, when standardized monitoring protocols were implemented. Nest entry timing was analysed using the full available record from 1998 to 2025. Breeding duration analyses were limited to 2012-2025 because reliable paired entry and exit dates were available only from 2012 onward.

For breeding duration, we included species-years with ≥ 2 nests to maximise temporal coverage while retaining sufficient within-year replication.

We assessed temporal trends using ordinary least-squares (OLS) linear regression with year as the predictor, reporting slopes, 95% CIs, p-values and R² after visually assessing model assumptions (Zuur et al. 2009).

For species-level nest occupancy and success models, we included nests monitored per species-year as a covariate to account for monitoring effort, and report the slope for Year.

To examine whether large-scale climate variability affected breeding, we used the continuous Multivariate ENSO Index (MEI) rather than classifying years into categorical ENSO states. Pre-breeding MEI was evaluated as an additional candidate predictor in the AICc model-selection framework described below. Associations between MEI and nest-entry timing were treated as exploratory because of the small number of species-year observations available.

#### 2.5.2 Climatic Drivers of Breeding Parameters

##### Model selection framework

We used information-theoretic model selection based on AICc (Burnham and Anderson 2002). For each breeding parameter, we constructed candidate linear models including temperature and rainfall from the entry, early breeding, and peak breeding windows as additive effects, together with a null (intercept-only) model. We ranked models by ΔAICc and considered ΔAICc < 2 to indicate substantial support, reporting AICc values, ΔAICc, weights, and parameter counts for each candidate set.

##### Phenological responses

Nest entry timing was analysed using a fixed-calendar sliding window approach with randomization control, described below. Annual nest occupancy and nesting success were modelled using fixed seasonal climate windows, with candidate models including temperature and rainfall from the entry, early breeding, and peak breeding windows, together with a null model; pre-breeding MEI was also included as a candidate predictor. We modelled breeding duration (mean days per species per year) as a function of the same seasonal climate variables, conducting analyses separately for each species.

##### Fixed-calendar sliding window analysis and randomization control

ing Sliding windows measured relative to each nest’s own entry date can be confounded with within-season temperature progression (Simmonds et al. 2019; van de Pol et al. 2016), therefore, we analysed nest-entry timing using an absolute sliding window analysis (SWA) design, in which the candidate window is anchored to a fixed calendar reference date rather than to the observed entry date. Reference dates were set *a priori* from known breeding biology, independent of observed entry dates: 1 March for Great and Wreathed Hornbill, 1 April for Oriental Pied Hornbill. We chose 1 March for Great and Wreathed Hornbills because it lies before the bulk of nest entries and before the long-term median entry dates (mid March for both species). The aim was to focus the candidate windows on conditions leading up to the typical entry period rather than on temperature during the main entry season itself.

For each species, we used the *climwin* package (van de Pol et al. 2016) to exhaustively test all candidate windows of mean daily temperature from 0 to 150 days before the reference date, ranking candidate models against an intercept-only baseline by AICc.

To guard against identifying a spuriously well-supported window from this exhaustive search, we ran a randomization control (climwin::randwin, 500 repeats per species): entry dates were shuffled and the full exhaustive window search was rerun on each shuffled dataset, generating a null distribution of best-achievable ΔAICc under no true relationship. Randomization p-values were calculated as the proportion of null-distribution values at least as extreme as the observed best window’s ΔAICc.

Because exhaustive search can occasionally select an isolated, non-credible window by chance, we applied a pre-specified credibility rule identically to all three species. The top-ranked window was retained unless it was spatially isolated in window position and length, defined as having fewer than three neighbouring windows within 10 days in window-open/window-close space that also scored within 2 ΔAICc units of the best value. If the top window was isolated, we selected the best-supported window within the largest cluster of alternative near-best windows.

Nest-years with more than 10% of their species’ 150-day candidate window overlapping a climate-station data gap (Section 2.4) were excluded rather than having missing temperature interpolated. We tested whether this exclusion introduced bias by comparing entry timing between retained and excluded nests (Wilcoxon rank-sum tests) and by comparing median entry timing across retained years versus all available years (Results).

We fit the selected window’s mean temperature in a linear mixed model with nest-entry day-of-year as the response, temperature and its interaction with species as fixed effects, and year as a random intercept (REML, Satterthwaite degrees of freedom; lme4/lmerTest). Because temperature varies at the species-year level rather than the individual-nest level, we additionally fit the relationship at the species-year level (median entry date per species per year) as a more conservative check on the level of independent replication.

To test whether the fitted relationship generalized beyond the years used to fit it, we conducted leave-one-year-out cross-validation: for each species, the model was refit after excluding one year, used to predict entry timing in that held-out year, mean absolute error (MAE) was calculated across held-out years and compared with the observed year-to-year variation in entry date as a baseline.

We used the same fixed-window/randomization framework to test rainfall as a candidate predictor. Where rainfall showed an association with entry timing, we assessed collinearity with the selected temperature window at the species-year level and compared nested models (temperature added to a rainfall-only baseline, and the reverse) to determine whether each variable explained independent variation.

As a supplementary check using no window-search procedure, we also fit a discrete-time hazard model of daily nest-entry probability as a function of a fixed 7-day trailing mean temperature. Because temperature and time-of-season are collinear within any single breeding season, we fitted models with and without an explicit control for within-season timing progression (day-of-season, modelled as a quadratic term). This analysis was treated as a supplementary diagnostic rather than as independent confirmation of the sliding-window result.

## Results

### 3.1 Long-term Trends in Breeding Parameters

#### Breeding phenology

Nest entry timing showed no significant change over time for Great Hornbill (slope = −0.005 days/year, 95% CI [−0.63, 0.62], p = 0.987, R² < 0.01, n = 21 years) or Wreathed Hornbill (slope = 0.08 days/year, 95% CI [−0.61, 0.76], p = 0.816, R² < 0.01, n = 22 years), using the full available monitoring record (1998-2025). Oriental Pied Hornbill, in contrast, showed a significant delay in entry timing over this period (slope = 0.90 days/year, 95% CI [0.10, 1.71], p = 0.031, R² = 0.29, n = 16 years; Figure 1, top row). Actual nesting occurred from mid-February onwards, with species-specific means and ranges reflecting consistent phenological windows: Great Hornbill (mean DOY 76.8 ± 12.1 SD, range: 17 February-19 April, n = 108 nests), Wreathed Hornbill (mean DOY 78.8 ± 13.9, range: 14 February-28 April, n = 91), and Oriental Pied Hornbill (mean DOY 95.4 ± 13.2, range: 7 March-4 May, n = 54).

**Figure 1.**
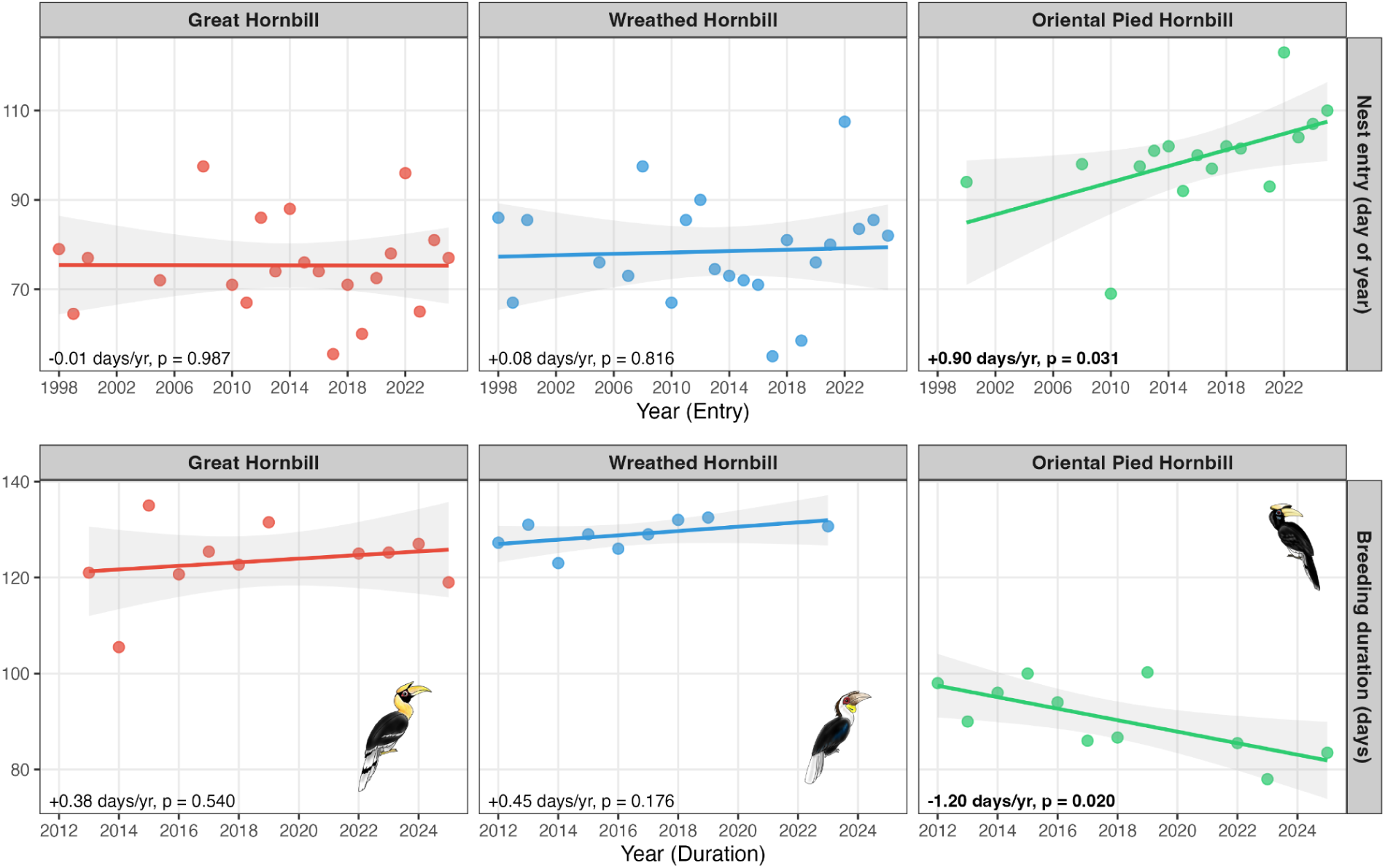
Long-term trends in nest entry timing (1998-2025, top row) and breeding duration (2012-2025, bottom row), by species. Points show annual values; lines show linear regression fits with 95% confidence intervals.

Breeding duration, analyzed for 2012–2025 (years with paired entry and exit dates), showed species-specific patterns (Figure 1, bottom row). Oriental Pied Hornbill breeding duration shortened significantly over time (slope = −1.20 days/year, 95% CI [−2.15, −0.24], p = 0.020, R² = 0.47, n = 11 years). Great Hornbill showed no significant change (slope = 0.38 days/year, 95% CI [−0.96, 1.71], p = 0.540, R² = 0.04, n = 11 years), and Wreathed Hornbill showed no significant change (slope = 0.45 days/year, 95% CI [−0.26, 1.16], p = 0.176, R² = 0.24, n = 9 years). Species differed substantially in mean breeding duration: Wreathed Hornbill (129 ± 3 days), Great Hornbill (123 ± 8 days), and Oriental Pied Hornbill (91 ± 7 days).

#### Population-level stability

Overall hornbill nest occupancy (all three species combined) showed no significant change over 27 years (1998-2025; slope = 0.22%/year, 95% CI [−0.36, 0.80], p = 0.442, R² = 0.02, n = 26 years; Figure S3A). Nesting success also showed no significant change (slope = −0.28%/year, 95% CI [−1.09, 0.53], p = 0.484, R² = 0.02, n = 22 years; Figure S3B). Both metrics exhibited substantial interannual variability, but no directional change was detectable over nearly three decades.

#### Species-specific nest occupancy trends

Nest occupancy varied among species during standardized monitoring (2007-2025), with mean annual occupancy of 77.4% (range 45-100%) in Great Hornbill, 71.7% (range 50-100%) in Wreathed Hornbill, and 70.7% (range 37.5-100%) in Oriental Pied Hornbill (Figure 2, top panel). Great Hornbill showed no significant temporal change in occupancy (slope = 1.46%/year, 95% CI [−0.30, 3.23], p = 0.098, R² = 0.17). Wreathed Hornbill showed a significant decline (slope = −1.56%/year, 95% CI [−2.74, −0.38], p = 0.013, R² = 0.33), whereas Oriental Pied Hornbill showed no significant change (slope = −0.60%/year, 95% CI [−2.21, 1.01], p = 0.443, R² = 0.27). Monitoring effort was not a significant predictor in any species model (p > 0.4 for all).

**Figure 2.**
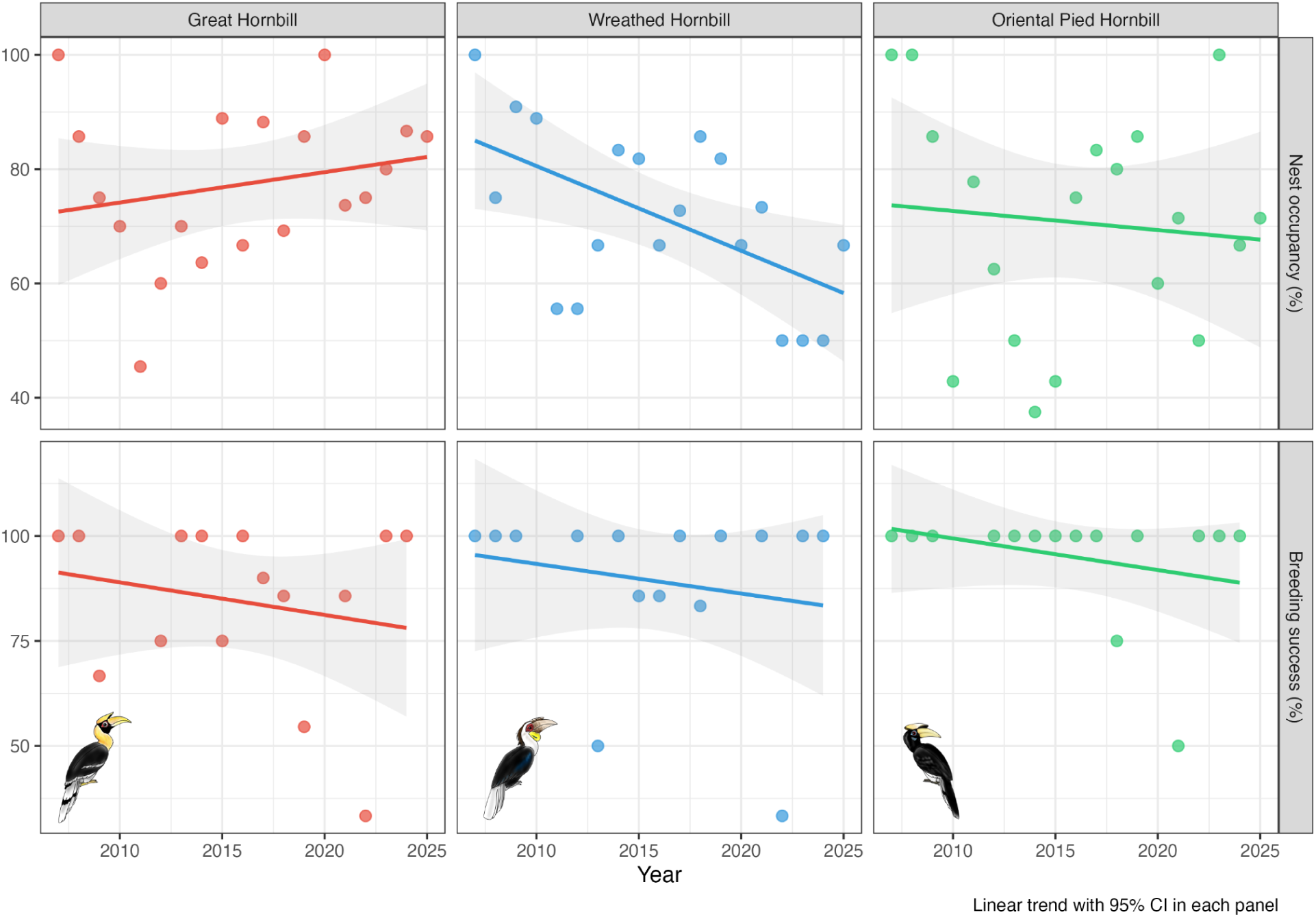
Species-specific patterns in breeding parameters (2007-2025). Nest occupancy during standardized monitoring is shown in the top panel. Wreathed Hornbill showed a significant decline; Great Hornbill and Oriental Pied Hornbill showed no significant trends. Lines show linear regressions with 95% confidence intervals. Nesting success in the bottom panel remained stable across all species. Points represent species-year values.

**Figure 3.**
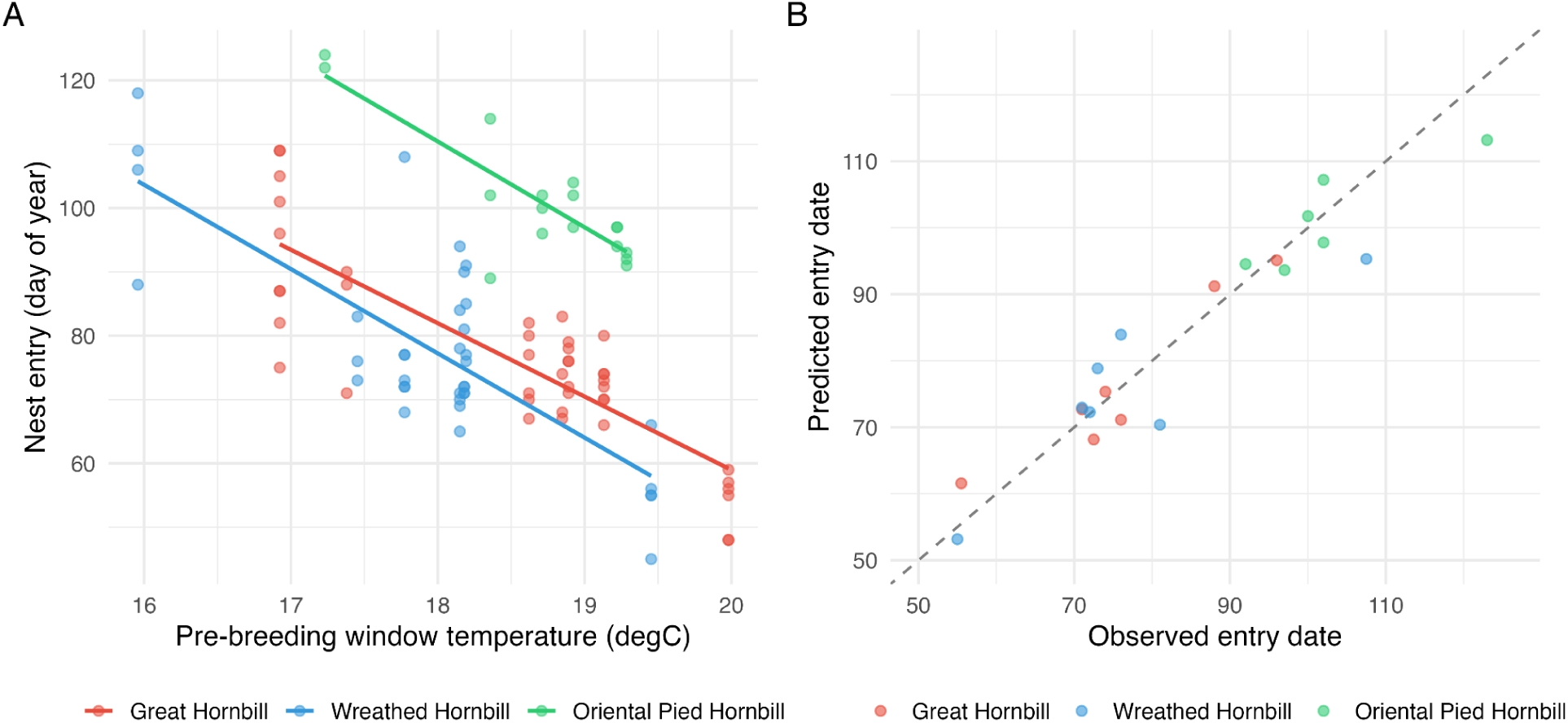
Interannual pre-breeding temperature predicts nest-entry timing, and this relationship generalizes to years not used to fit the model. (A) Nest-entry day-of-year against mean temperature in each species’ selected fixed-calendar window (Table 1); lines show fitted values from the joint mixed model. (B) Leave-one-year-out cross-validated predictions: each point shows a species-year predicted entry date from a model refit with that year excluded, plotted against the observed species-year median entry date; the dashed diagonal indicates perfect prediction.

**Table 1.** Fixed-calendar absolute sliding-window results for nest-entry timing, per species, with randomization control. Windows are expressed as calendar-date intervals relative to the species-specific reference date. The AICc difference is calculated as AICc of the selected climate model minus AICc of the intercept-only model; negative values indicate better support for the climate model than for the null model. Randomization *p* values are based on 500 exhaustive-search randomizations. Confidence intervals represent standard estimation uncertainty for the selected temperature window.

| Species | n (years) | Selected window | Slope (days per degC) | 95% CI lower | 95% CI upper | AICc difference from null | Randomization p |
| --- | --- | --- | --- | --- | --- | --- | --- |
| Great Hornbill | 7 | 9-15 February | -11.58 | -13.93 | -9.23 | -48.27 | 0.008 |
| Wreathed Hornbill | 7 | 26 January-14 February | -13.38 | -17.01 | -9.74 | -31.04 | 0.016 |
| Oriental Pied Hornbill | 6 | 12 January-4 March | -13.36 | -17.63 | -9.09 | -18.37 | 0.154 |

#### Nesting success by species

Nesting success remained stable across all three species during 2007-2025 (Figure 2, bottom panel), controlling for monitoring effort. Great Hornbill showed no significant change (slope = −1.06%/year, 95% CI [−4.67, 2.55], p = 0.533, R² = 0.05). Wreathed Hornbill showed no significant change (slope = −1.00%/year, 95% CI [−3.29, 1.28], p = 0.358, R² = 0.13). Oriental Pied Hornbill showed no significant change (slope = −0.80%/year, 95% CI [−2.34, 0.73], p = 0.278, R² = 0.12).

#### Local climate variability during the study period

Local climate over the study period (2011-2025) showed substantial interannual variation with no clear directional trends (see Fig. S3). Trend models were fitted only to years meeting a ≥80% climate data-coverage threshold for the relevant period; the resulting sample sizes were small for several sub-seasonal windows (n = 4-9 years) and these trends should be interpreted cautiously. Annual mean temperature did not change significantly over time (slope = 1.7°C/decade, 95% CI [−1.1, 4.5], p = 0.12, n = 4 years), and total annual rainfall likewise showed no significant trend (slope = −160 mm/decade, 95% CI [−1258, 938], p = 0.67, n = 5 years). Pre-breeding mean temperature ranged from 16.9 to 19.9°C (mean = 18.7 ± 0.9°C SD) and pre-breeding rainfall from 0.5 to 68 mm (mean = 31.1 ± 22.9 mm SD), with no significant trends in either variable (temperature slope = −0.3°C/decade, p = 0.78, n = 9; rainfall slope = 5 mm/decade, p = 0.82, n = 9). Breeding-season climate showed no significant trend in mean temperature (slope = 0.6°C/decade, p = 0.23, n = 7) and a marginal, non-significant decline in rainfall (slope = −1533 mm/decade, 95% CI [−3220, 153], p = 0.067, n = 7).

#### ENSO effects

Continuous pre-breeding Multivariate ENSO Index (MEI) values did not improve prediction of nest occupancy or nesting success beyond the null model in AICc model selection (occupancy: MEI model AICc weight = 0.007 vs. null = 0.993; success: MEI weight = 0.016 vs. null = 0.984; Table S1), indicating no detectable effect of large-scale climate oscillations on these breeding outcomes. Pre-breeding MEI was weakly and non-significantly correlated with pre-breeding temperature (r = 0.34), suggesting that pre-breeding MEI and local pre-breeding temperature captured relatively distinct aspects of interannual climate variation at the site. At the species-year level, pre-breeding MEI showed a consistent, negative, but non-significant association with nest-entry timing in all three species (Great Hornbill: r = −0.26, p = 0.57; Wreathed Hornbill: r = −0.42, p = 0.35; Oriental Pied Hornbill: r = −0.47, p = 0.34).

### 3.2 Climatic Drivers of Breeding Parameters

#### Entry timing and climatic associations

Excluding nest-years with substantial climate-data gaps (Methods; Table S2) retained 95 of 195 available nest-entry events across seven breeding seasons (2014-2018, 2020, and 2022; Great Hornbill *n* = 42, Wreathed Hornbill *n* = 36, Oriental Pied Hornbill *n* = 17). Excluded nests did not differ significantly in entry timing from retained nests in any species (Wilcoxon tests: Great Hornbill, *p* = 0.485; Wreathed Hornbill, *p* = 0.303; Oriental Pied Hornbill, *p* = 0.725). Applying the pre-specified credibility rule retained the top-ranked window for Great Hornbill and Oriental Pied Hornbill. For Wreathed Hornbill, the raw top-ranked window was an isolated candidate and was therefore replaced by the best-supported alternative under the pre-specified rule. Across the 500 randomization repeats, 408, 340, and 265 distinct window combinations were selected as best for Great Hornbill, Wreathed Hornbill, and Oriental Pied Hornbill, respectively, confirming that the exhaustive search was rerun for each randomized dataset.

Using the fixed-calendar absolute sliding-window design, pre-breeding temperature was a predictor of nest-entry timing in all three species. For Great Hornbill, the selected window was 9-15 February, corresponding to 20-14 days before the 1 March reference date; the slope was −11.58 days °C⁻¹ (95% CI: −13.93 to −9.23; ΔAICc = −48.27; randomization *p* = 0.008). For Wreathed Hornbill, the selected window was 26 January-14 February, corresponding to 34-15 days before the 1 March reference date; the slope was −13.38 days °C⁻¹ (95% CI: −17.01 to −9.74; ΔAICc = −31.04; randomization *p* = 0.016). For Oriental Pied Hornbill, the selected window was 12 January-4 March, corresponding to 79-28 days before the 1 April reference date; the slope was −13.36 days °C⁻¹ (95% CI: −17.63 to −9.09; ΔAICc = −18.37; randomization *p* = 0.154).

Fitting all three species jointly in a linear mixed model (*n* = 95 nests, 7 years), the temperature effect was highly significant (β = −11.50 days °C⁻¹, *p* < 0.0001). Neither species interaction term was significant (Wreathed Hornbill: β = −1.71, *p* = 0.400; Oriental Pied Hornbill: β = −1.95, *p* = 0.580), indicating that the data do not provide strong statistical evidence that temperature sensitivity differs among species despite differences in point estimates. Species-year analyses showed negative temperature associations for Great Hornbill (slope = −12.0 days °C⁻¹, *p* = 0.00017), Wreathed Hornbill (slope = −14.2 days °C⁻¹, *p* = 0.0014), and Oriental Pied Hornbill (slope = −13.4 days °C⁻¹, *p* = 0.0022). Leave-one-year-out cross-validation showed prediction error (MAE) of 3.2, 5.8, and 4.5 days for Great, Wreathed, and Oriental Pied Hornbills, respectively, smaller in each case than the species’ observed year-to-year standard deviation in entry date (12.9, 15.8, and 10.7 days). These results indicate that the fitted relationship generalized to left-out years, although the number of species-years was small.

Rainfall during the same pre-breeding period showed a substantial univariate association with entry timing in Great Hornbill and Wreathed Hornbill, but rainfall was correlated with the selected temperature window in both species (*r* = 0.79 and 0.73, respectively). Temperature retained significant explanatory power when added to a rainfall-only model in Great Hornbill (*F* = 28.56, *p* = 0.0059) and Wreathed Hornbill (*F* = 13.35, *p* = 0.0217), whereas rainfall did not improve fit when added to a temperature-only model in either species (Great Hornbill: *F* = 0.81, *p* = 0.418; Wreathed Hornbill: *F* = 1.30, *p* = 0.317). We therefore treat temperature as the better-supported predictor of nest-entry timing in these two species. For Oriental Pied Hornbill, only five complete species-years were available for the temperature–rainfall comparison; neither temperature nor rainfall provided evidence of an independent contribution in the nested models (temperature added to rainfall-only: *F* = 2.30, *p* = 0.269; rainfall added to temperature-only: *F* = 0.51, *p* = 0.548). The Oriental Pied Hornbill relation to rainfall is therefore considered unresolved rather than as a confirmed independent predictor.

A discrete-time hazard model, testing whether daily temperature predicted the instantaneous probability of nest entry without any window-search procedure, initially showed a positive association (temperature coefficient = 0.48). However, this association did not hold when a control for within-season timing progression was added (coefficient = −0.09, *p* = 0.337), reflecting strong collinearity between recent temperature and time of season during the period when entries occur. We therefore do not treat the hazard model as independent confirmation of the fixed-window result.

#### Species-specific breeding duration responses

Breeding duration showed no support for any climate window over the null model in any species (Table S3). The null model was the best-supported model for Great Hornbill, Wreathed Hornbill, and Oriental Pied Hornbill, with every climate-window alternative more than 34 AICc units higher than the null model. Because no non-null model had ΔAICc < 2, no climate-window parameter estimates are reported as supported effects. We therefore find no evidence that breeding duration was predicted by climate at the species-year scale.

#### Annual breeding outcomes show no evidence of climate association

In contrast to the phenological response to temperature, annual nest occupancy and nesting success showed no evidence of being predicted by interannual climate variation. All occupancy climate models ranked far below the null model (ΔAICc > 33 for every candidate), and nesting-success climate models likewise provided no explanatory support (ΔAICc > 25 for every candidate). This decoupling indicates that, while the timing of breeding tracks interannual temperature variation in all three species, annual breeding outcomes show no detectable climate sensitivity at this site, consistent with the long-term stability observed in nest occupancy and nesting success trends.

## Discussion

We addressed whether tropical hornbill breeding tracks interannual climate variability or long-term trends, and which climatic variables best explain interannual variation in nest-entry timing. Pooled nest occupancy and nesting success showed no directional change over the long-term monitoring period despite ∼1.2 °C regional warming, and continuous pre-breeding MEI did not improve prediction of occupancy or success, indicating no detectable population-level response of these breeding outcomes to large-scale climate variability. Interannual pre-breeding temperature was a predictor of nest-entry timing in all three species, with warmer conditions associated with earlier entry, although randomization support was stronger for Great and Wreathed Hornbills than for Oriental Pied Hornbills.. Temperature retained explanatory power over rainfall in Great and Wreathed Hornbills, whereas rainfall did not add independent explanatory power once temperature was accounted for; the temperature-rainfall relationship could not be cleanly resolved for Oriental Pied Hornbill. Despite this interannual climatic sensitivity, long-term nest-entry timing remained stable in Great and Wreathed Hornbills, while Oriental Pied Hornbill showed a significant delay in entry timing and shortening of breeding duration, and Wreathed Hornbill showed a significant decline in nest occupancy. These patterns indicate that interannual phenological sensitivity does not necessarily translate into directional long-term change or detectable changes in annual reproductive outcomes.

### Long-term stability despite environmental change

The absence of directional change in pooled nest occupancy and nesting success, together with stable nest-entry timing in Great and Wreathed Hornbills, suggests substantial buffering of breeding phenology and reproductive performance despite long-term environmental change. This contrasts with many temperate passerines, which have advanced laying dates by roughly 2-5 days per decade in response to warming spring temperatures (e.g. Dunn & Winkler 1999). At Pakke, local climate records showed no significant directional trend in pre-breeding temperature or rainfall over the years with adequate data coverage, while nest-entry timing varied substantially among years. This combination suggests that hornbills may be responsive to interannual climate variation without exhibiting a simple progressive shift in mean breeding timing.

Three features may contribute to this pattern. First, the absence of a detectable trend in local pre-breeding temperature means that the climatic variable most strongly associated with nest-entry timing has itself not shown a sustained directional change in the available local record. Second, substantial interannual variation in nest-entry timing provides scope for flexible adjustment among years without necessarily producing a long-term trend. Third, phenological plasticity may allow hornbills to adjust breeding timing within a relatively constrained seasonal window while maintaining breeding performance.

This pattern is consistent with the broader expectation that tropical climate change may register ecologically through variability rather than linear trends (Trisos et al. 2020; Vasseur et al. 2014). Where temperate phenological studies often document progressive directional shifts over decades (e.g. Dunn & Winkler 1999), our results show that tropical hornbills can exhibit strong sensitivity to year-to-year temperature variation while showing little directional change in pooled breeding outcomes or, for Great and Wreathed Hornbills, in mean nest-entry timing. Thus, a framework focused only on long-term trends could overlook substantial within-system climatic sensitivity. At the same time, the significant delay in Oriental Pied Hornbill nest entry and shortening of its breeding duration, together with the decline in Wreathed Hornbill nest occupancy, indicate that stability is not uniform across species and that species-specific changes warrant continued attention. This distinction may be broadly relevant to tropical avian phenology: species living close to historically narrow climatic envelopes (Deutsch et al. 2008) may show their clearest climate responses not as a trend line, but as sensitivity to the magnitude of each year’s departure from the local norm.

The species-specific long-term patterns suggest different dimensions of change rather than a uniform response across hornbills. Wreathed Hornbill showed a significant decline in nest occupancy of 1.56 percentage points per year during standardized monitoring, while breeding success remained stable. One possible explanation is increasing competition for a limited supply of suitable nest cavities. Because the three hornbill species differ substantially in body size and cavity-use requirements, competition may be asymmetric, with larger species potentially displacing smaller species from overlapping nest sites. Interspecific competition and nest-site takeovers have been documented among hornbills at Pakke, including takeovers involving Great and Wreathed Hornbills (Datta & Rawat 2004).

Great Hornbill showed no significant change in either occupancy or success, although occupancy showed a non-significant positive tendency. Oriental Pied Hornbill showed no significant change in occupancy or success, but nest entry became significantly later by 0.90 days per year and breeding duration shortened by 1.20 days per year. These directional changes were not explained by the climate variables tested: local pre-breeding temperature showed no significant long-term trend, annual occupancy and nesting success were not supported by climate models, and breeding duration showed no support for any climate window. The species-specific trends therefore warrant continued monitoring and investigation of other ecological processes, including changes in food availability, cavity availability or competition, rather than as evidence of a climate-driven response.

### Temperature, not rainfall, predicts phenological variation

Temperature was the better-supported climatic predictor of hornbill phenology, though rainfall was not unrelated to breeding timing. Pre-breeding rainfall showed an association with entry timing in Great and Wreathed Hornbills, but this association was not independent of temperature. The two variables were strongly correlated during this period, temperature retained explanatory power when rainfall was included in the same model, and rainfall did not improve model fit once temperature was accounted for. For Oriental Pied Hornbill, temperature and rainfall were very strongly negatively correlated, preventing clear separation of their effects. This pattern is consistent with theoretical expectations that organisms in seasonal environments track reliable predictors of resource availability (Burnside et al. 2021), and suggests that temperature and rainfall may co-vary as part of a shared seasonal climate signal at this site, plausibly linked through their joint influence on flowering and fruiting phenology (Datta et al. 2025), rather than acting as fully independent cues.

The negative relationships between pre-breeding temperature and nest entry indicate that hornbills track interannual temperature variation in the timing of breeding, though we cannot determine whether temperature itself is directly perceived or is a correlated proxy for an underlying resource signal. The record also included years with unusually early nest entry, illustrating that the response was not simply a shift toward delayed breeding under cooler conditions. Pre-breeding MEI was only weakly correlated with the temperature signal and showed a consistent but non-significant negative association with entry timing in all three species. The latest entry in all three species occurred in 2022, during a period in which La Niña conditions persisted across several consecutive years (2021-2023). This is consistent with, but cannot confirm, an ENSO contribution to the same interannual signal.

### Temperature sensitivity is similar across species

Point estimates of temperature sensitivity were similar across species (Great Hornbill: −12.0; Wreathed Hornbill: −14.2; Oriental Pied Hornbill: −13.4 days per °C), and the temperature × species interaction was not statistically significant in the joint model. The data therefore provide no evidence that sensitivity differs among species, although with seven years of climate data the power to detect such differences is low. Because temperature varies at the species-year rather than individual-nest level, the effective replication for this relationship is the number of species-years rather than the number of nests. The species-year analysis therefore provides the more conservative basis for interpreting the temperature association.

### Downstream effects on breeding outcomes

Pre-breeding temperature predicted nest timing in all three species, but downstream breeding parameters showed no evidence of comparable climate sensitivity. Breeding duration was not predicted by any climate window in any species (null model favoured in all three), and annual occupancy and success were unrelated to temperature, rainfall or MEI (all climate models ranked well below the null, ΔAICc > 25).

This decoupling is consistent with phenological plasticity buffering populations from potential reproductive consequences of climate variability, though failure to detect a climate effect on occupancy or success is not itself direct evidence of a buffering mechanism, particularly given the few years available for the climate analyses. The interannual record also shows years where this apparent buffering may have been compromised. Early nesting in the warm year of 2017 occurred without evident downstream consequences, whereas 2019 and 2022 represented contrasting anomalous years: in 2019, nest entry was relatively early in Great Hornbills (day 60) and Wreathed Hornbills (day 58.5), while nesting success was 54.55% in Great Hornbills and 100% in both Wreathed and Oriental Pied Hornbills; in 2022, nest entry was substantially later in all three species—day 96, 107.5, and 123, respectively—and nesting success declined to 33.33% in Great and Wreathed Hornbills, although it remained 100% in Oriental Pied Hornbills. The 2022 pattern coincided with a cooler entry-period temperature than in 2019 (16.90°C versus 18.68°C) and a multi-year La Niña episode (2021 to 2023). Based on these individual years, this suggests but does not show that extreme departures may exceed the range within which timing shifts alone can maintain reproductive performance. Whether phenological tracking of temperature maintains synchrony with fruiting phenology and secures provisioning resources, as opposed to acting through some other pathway, remains to be explored.

### Comparative context: African hornbills and tropical-temperate differences

Our findings contrast with studies of some African hornbill species showing strong negative effects of temperature and rainfall on breeding success (van de Ven et al. 2020; Pattinson et al. 2022). This divergence may reflect differences in climatic regime rather than temperature alone. Pakke experiences high temperatures (max ∼40°C) during the breeding season, but these occur within a humid, high-rainfall monsoon system, whereas the African study systems are in arid zones characterized by hotter and drier conditions and greater rainfall variability. The use of nest boxes in the African studies may also contribute to differences in thermal exposure relative to natural cavities, which provide different microclimatic conditions and show stronger negative impacts of climate (Combrink et al. 2017; Pattinson et al. 2022). More generally, if tropical frugivores and their food plants respond to the same seasonal climate cues, breeding may remain more closely synchronized with resource availability than in systems where climate-driven shifts in breeding and food phenology become decoupled.

### Implications and future directions

All three hornbill species tracked interannual pre-breeding temperature in their nest-entry timing, while pooled nest occupancy and breeding success showed no long-term directional change. At the same time, the species-specific results show that this overall stability was not uniform: Wreathed Hornbill occupancy declined, and Oriental Pied Hornbill showed later nest entry and shorter breeding duration. These latter changes were not explained by the climate variables tested. Together, the results suggest that tropical hornbill breeding can be highly responsive to year-to-year temperature variation without necessarily producing a corresponding directional change in reproductive performance.

The observed temperature sensitivity in nest entry timing, nevertheless needs to be interpreted within the climatic range and temporal replication of this study. The climate analyses rest on seven breeding seasons (2014-2018, 2020, 2022) because multi-month station outages removed the pre-breeding window in other years, and 2024 and 2025 could not be added for the same reason. Temperature varies at the species-year level, so the effective replication for the temperature effect is seven years rather than the number of nests. Oriental Pied Hornbill has the smallest effective sample (six years), which limits power to distinguish its temperature effect from alternative explanations. The Wreathed Hornbill window was a single alternative candidate after its top-ranked window was rejected, so it should be interpreted more cautiously than the Great Hornbill window. Finally, temperature is a predictor here, not an established cue: we cannot determine whether the association with nest-entry timing reflects direct thermal effects or a correlated signal of environmental conditions that influence the resources on which hornbills depend.

Projected warming therefore should not be translated directly into an expected number of days of phenological advance. The observed slopes describe responses within the range of interannual conditions represented in the climate-analysis years, and it is unknown whether these relationships remain linear under warming beyond that range or whether earlier breeding would remain synchronized with fruiting phenology. High ambient humidity during the breeding season adds a further concern: as humidity rises, evaporative cooling becomes impaired, and wet-bulb temperature rather than air temperature becomes the ecologically relevant measure of heat stress (Nord et al. 2026; Coulson et al. 2025). Birds breeding in wet tropical forests may therefore face a compound temperature–humidity threat that is underestimated by air-temperature projections alone.

The observed temperature sensitivity of nest-entry timing raises an important question about what ultimately constrains breeding in these hornbills. If temperature acts partly through environmental conditions that determine food availability, then phenological tracking could help maintain synchrony between breeding and resource availability under interannual climatic variation. Alternatively, temperature may act through direct physiological or behavioural effects on breeding decisions. Distinguishing these possibilities will require concurrent measurements of climate, fruiting phenology, and hornbill breeding across more years and populations. Such data would also help determine whether the observed plasticity can persist under warming beyond the range experienced during our study, and whether the directional changes observed in Oriental Pied Hornbill and Wreathed Hornbill reflect climate sensitivity or other ecological processes.

## ACKNOWLEDGEMENTS

We thank the Arunachal Pradesh Forest Department for research permits and support, especially Pekyom Ringu, G.N. Sinha, Bharat Bhatt, N. Tam, Dr. Damodhar A.T., Millo Tasser, and Tajum Yomcha. The Pakke Tiger Reserve management facilitated fieldwork over many years; we particularly thank Tana Tapi, Suraj Singh, Satyaprakash Singh, Dhawan Kumar Rawat, P.B. Rana, Rubu Tado, Kime Rambia, Singku Maga, and the wider frontline staff of the reserve. The field technicians of the Eastern Himalaya Programme, NCF, were indispensable to long-term data collection: the late Kumar Thapa, late Tali Nabum, late Narayan Mogar, Peter Wage, Jacob Brah, Sagar Kino, Arjun Rai, Sital Dako, and Rasham Brah. Research coordination across seasons/years was provided by Amruta Rane, Ushma Shukla, Devathi Parashuram, Bibidishananda Basu, Noopur Borawake, Akanksha Rathore, Swati Sidhu, Saniya Chaplod, and Sana Huque. Rom Whitaker and the Agumbe Rainforest Research Station donated the weather station in 2011; Shankar CM and Kamal Sonar provided technical support. Funding was provided by the National Geographic Society, Disney Wildlife Conservation Fund, Wings Women of Discovery, Wildlife Conservation Society, Whitley Fund for Nature, Stop Poaching Fund, The Serenity Trust, and a British Ecological Society grant. We are thankful to Arjun Srivathsa for the hornbill illustrations.

## AUTHOR CONTRIBUTIONS

AD conceived, initiated and acquired funding for this long-term project. AD, RN, KP, KT, JB participated in field data collection along with other members in the field team over the study period. Data curation was done by AD, KP, RN, SB and RC. RC completed formal data analyses with contributions from AD, SB and RN. RC and AD led the writing of the paper with inputs from RN, SB and KP.

