## Supplementary material for "Tracking interannual temperature variation: phenological plasticity and reproductive stability in tropical hornbills": Supp Figures and Tables

**Supplementary Figures and Tables**

**Weather Patterns**

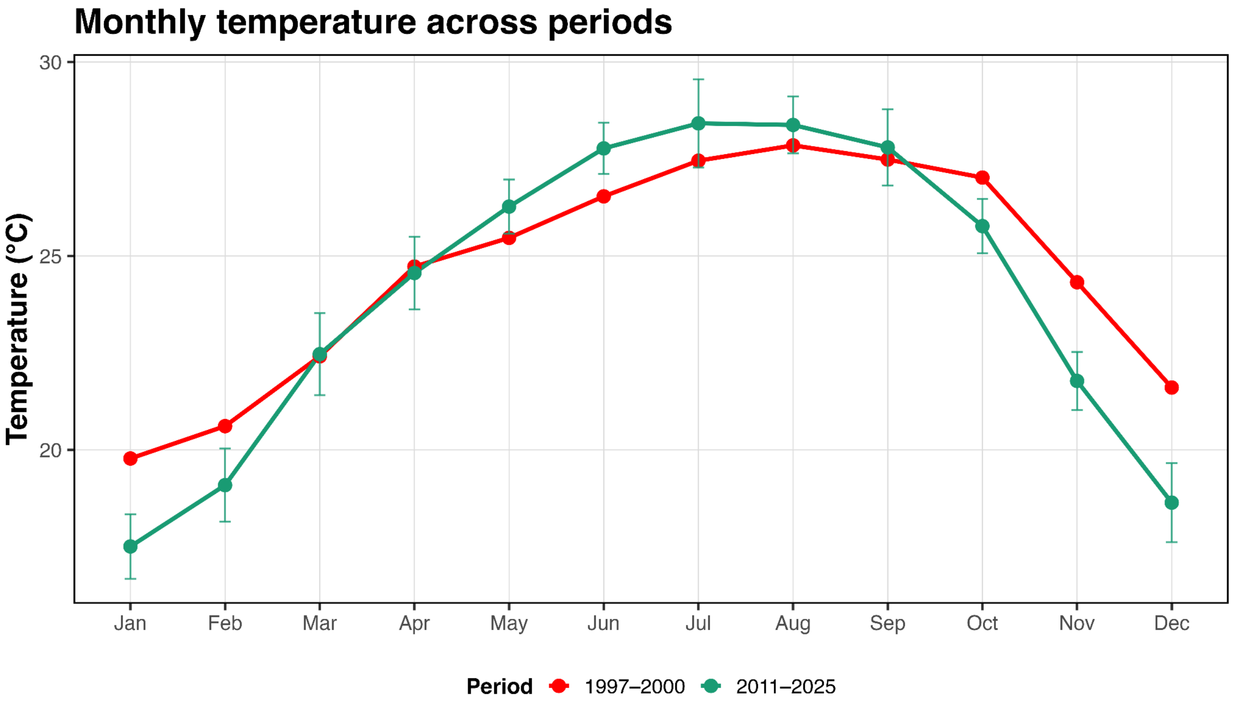


**Figure S1.** Monthly temperature across periods, showing the 1997-2000 baseline (red) and the 2011-2025 (green) study period. Lines show monthly mean temperature, and error bars show interannual variability for 2011-2025, allowing comparison of the seasonal baseline shift between the two periods.

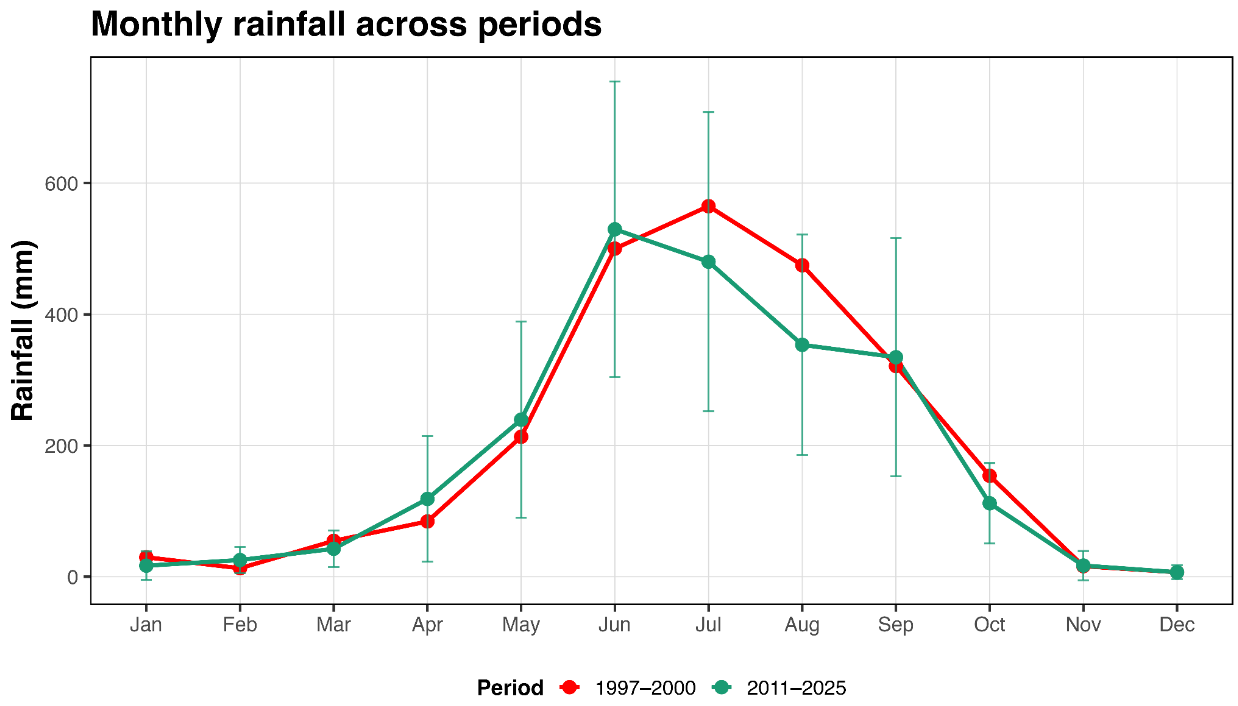


**Figure S2.** Monthly rainfall across periods, showing the 1997-2000 baseline (red) and the 2011-2025 (green) study period. Lines show mean monthly rainfall, and error bars show interannual variability for 2011-2025, allowing comparison of the seasonal rainfall pattern and baseline shift between the two periods.

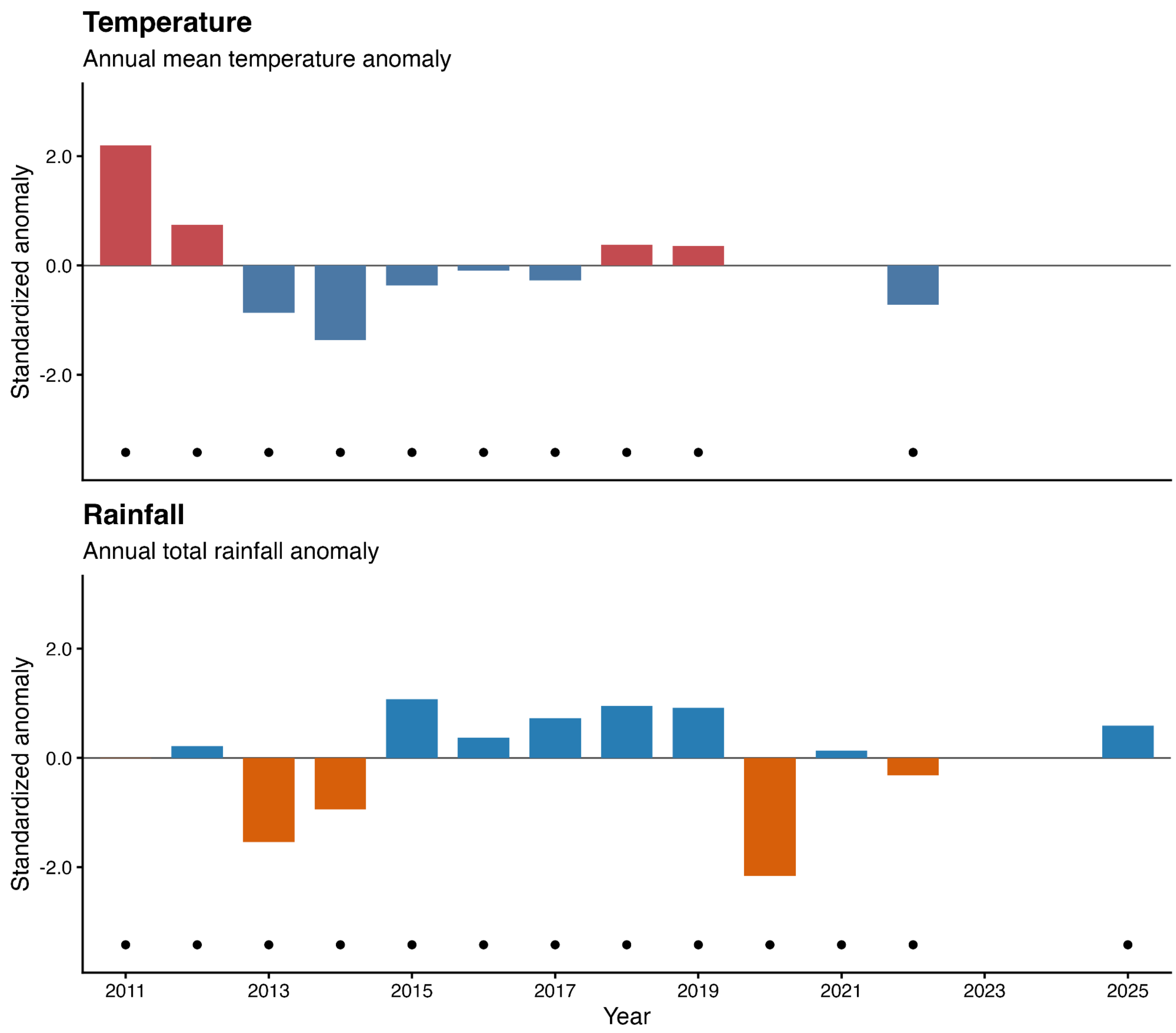


**Figure S3. Interannual variability in local temperature and rainfall at Pakke, 2011–2025.** Bars show standardized annual anomalies calculated from quality-filtered local weather-station records. Positive and negative values indicate warmer/cooler or wetter/drier years, respectively, relative to the mean of adequately sampled years for the corresponding variable. Black points below the x-axis indicate years meeting the 80% data-coverage threshold. Years below this threshold remain on the time axis but are not used to calculate or plot anomalies. The figure is descriptive and is intended to show year-to-year climatic variation, not to estimate a long-term climate trend.

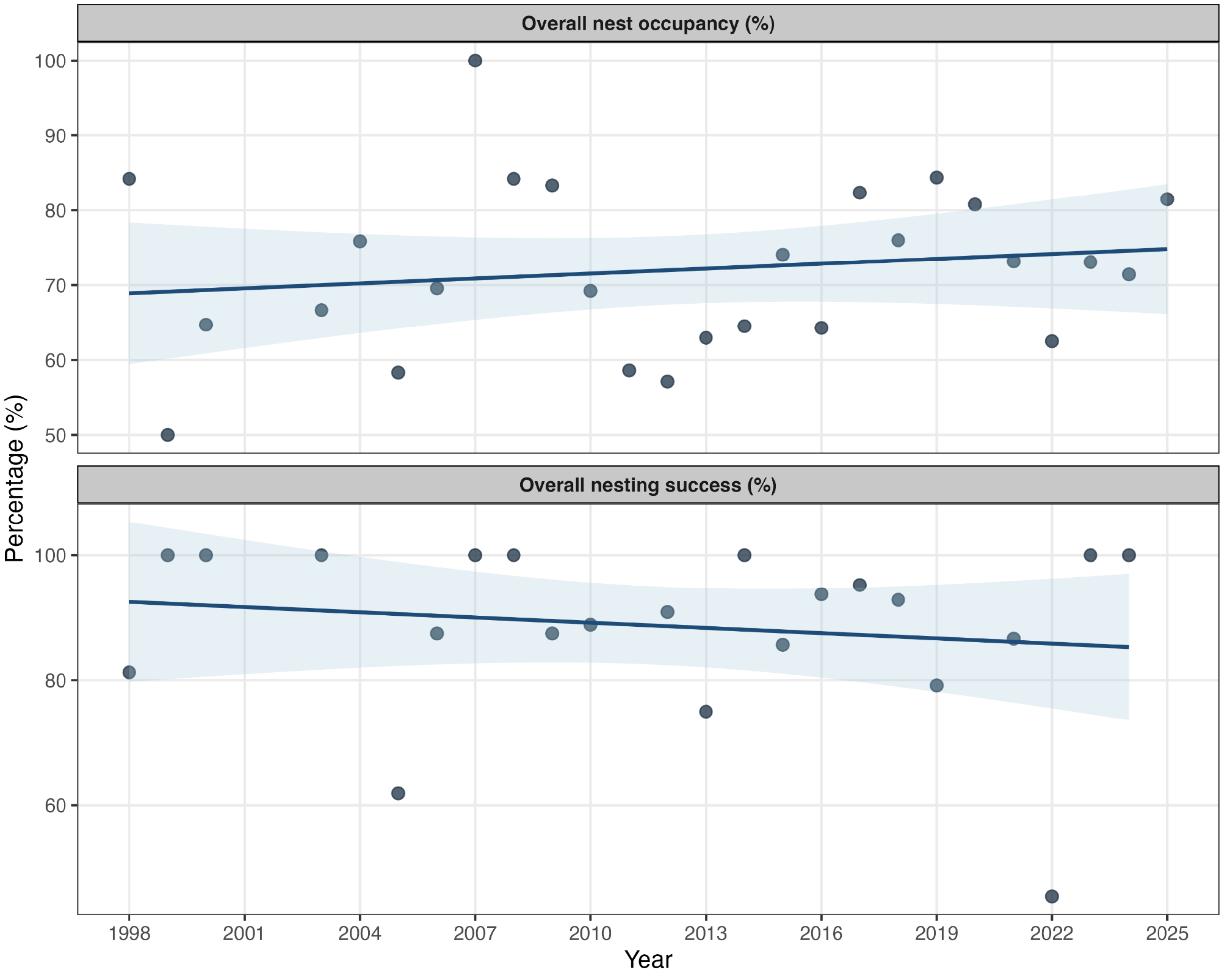


**Figure S4. Long-term trends in overall hornbill breeding performance, 1998-2025.** (A) Overall nest occupancy showed no significant temporal trend from 1998 to 2025 (slope = 0.22 percentage points/year, 95% CI -0.36 to 0.80, $p = 0.442$, $R^2 = 0.025$). (B) Overall nesting success also showed no significant temporal trend (slope = -0.28 percentage points/year, 95% CI -1.09 to 0.53, $p = 0.484$, $R^2 = 0.025$). Points represent annual percentages; lines show ordinary least-squares linear regressions with shaded 95% confidence intervals. Missing values indicate years for which the relevant parameter was unavailable.

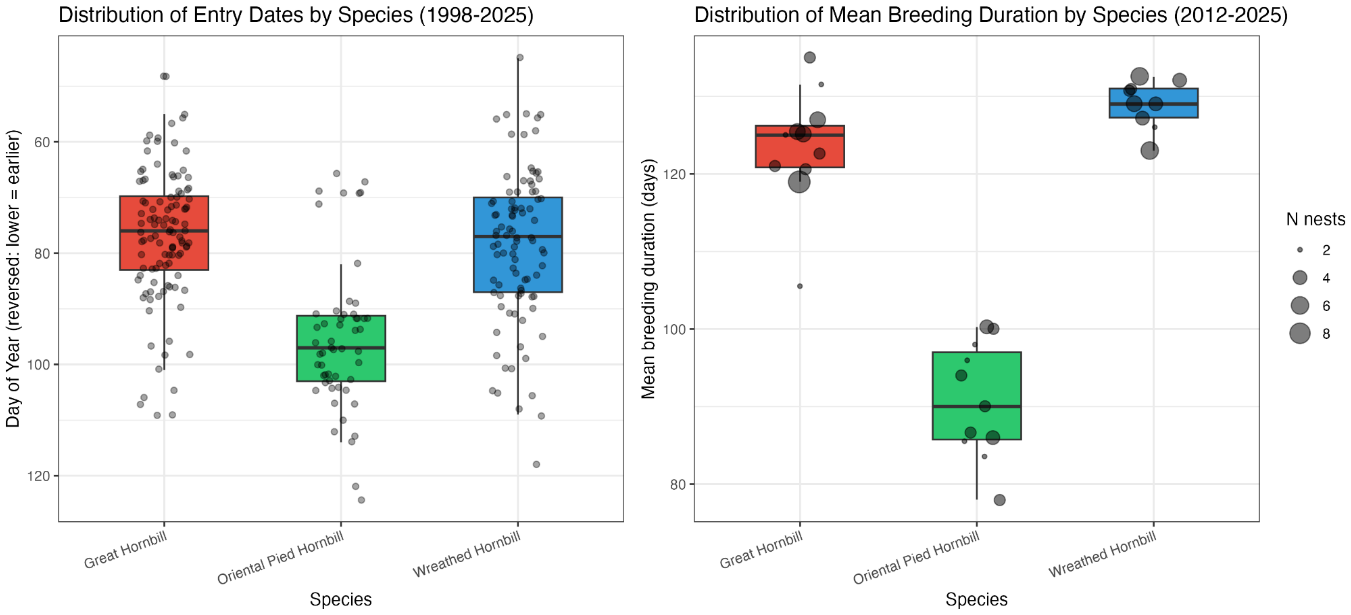


**Figure S5. Distribution of nest-entry timing and breeding duration by hornbill species.** (A) Individual nest-entry dates (day of year) for the full monitoring record, 1998-2025. (B) Mean breeding duration for each species-year, 2012-2025. Breeding duration could only be calculated for nests with both reliable entry and exit dates, which were not available for every nest, and species-years with fewer than two such nests were excluded; the number of observations therefore varies among species and years. Points show individual entry dates (A) or species-year means sized by the number of nests (B). Boxplots show the median, interquartile range, and range.

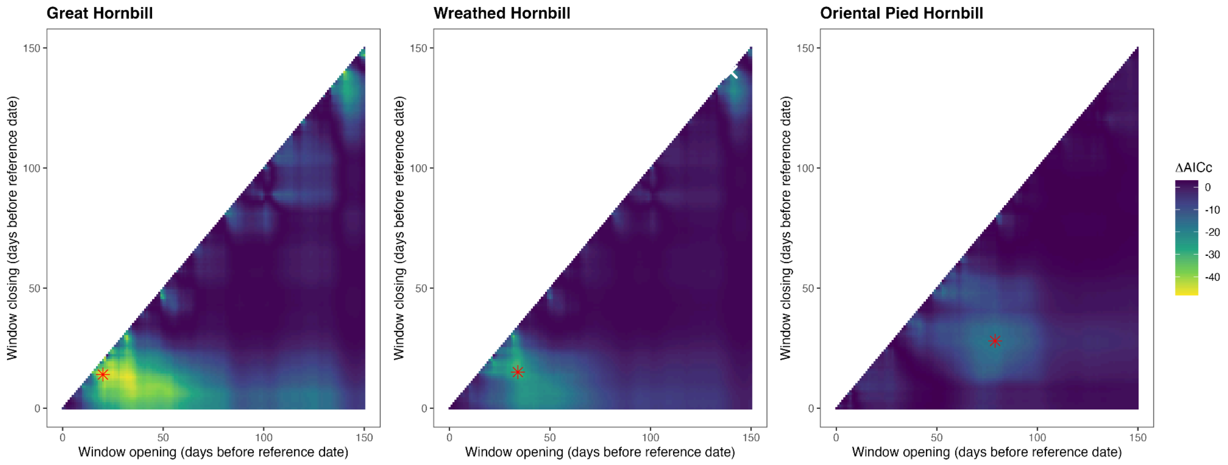


**Figure S6. Support for candidate temperature windows associated with nest-entry timing. Each square is one candidate window.** The horizontal axis shows how many days before the species' fixed reference date (1 March for Great and Wreathed Hornbill; 1 April for Oriental Pied Hornbill) the window opens, and the vertical axis shows how many days before that date it closes. A window cannot close before it opens, so only the area below the diagonal is filled. Windows between 0 and 150 days before the reference date were tested. Colour shows ΔAICc, the improvement in fit relative to a model without temperature: more negative values (yellow) mean that mean temperature in that window predicts entry date better. The red star marks the window retained after applying the pre-specified credibility rule. In the Wreathed Hornbill panel, the white × marks the top-ranked window, which was rejected because it was isolated (no similarly supported windows nearby), and the red star marks the best alternative. The selected windows were 20-14 days before 1 March for Great Hornbill (9-15 February), 34-15 days before 1 March for Wreathed Hornbill (26 January-14 February), and 79-28 days before 1 April for Oriental Pied Hornbill (12 January-4 March).

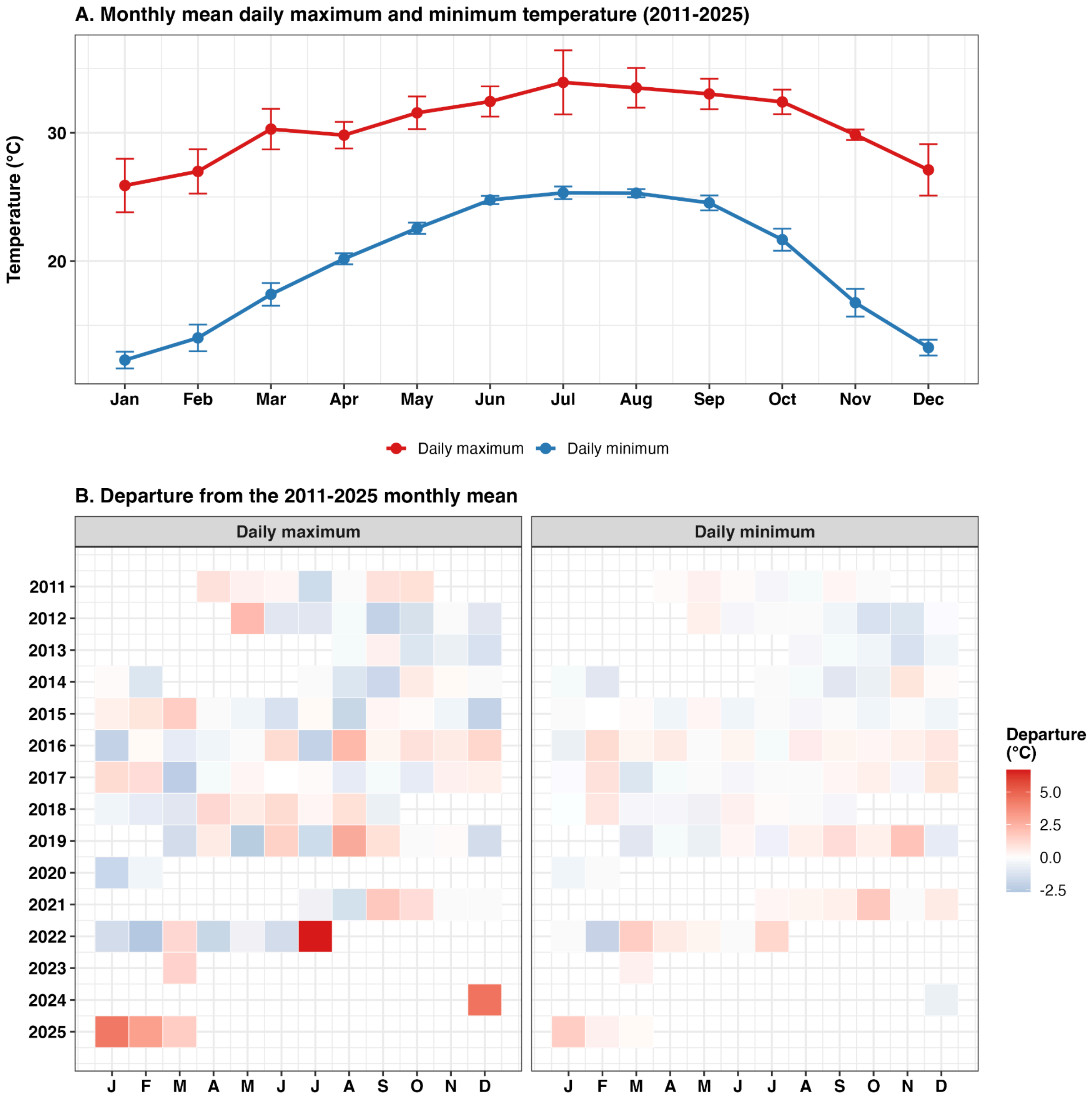


**Figure S7: Monthly daily maximum and minimum temperatures at the study site, 2011-2025.** (A) Monthly means of daily maximum and minimum temperatures, with error bars showing interannual standard deviations. (B) Differences between each month's mean daily maximum and minimum temperatures and the corresponding 2011-2025 averages. Months with fewer than 20 days of valid data are left blank.

| **Response** | **Model** | ***K*** | **AICc** | **ΔAICc** | **AICc weight** |
|---|---|---|---|---|---|
| Nest occupancy | Pre-breeding MEI | 3 | 60.21 | 9.86 | 0.007 |
| Nest occupancy | Entry temperature + rainfall | 4 | 84.07 | 33.73 | 0.001 |
| Nest occupancy | Early-season temperature + rainfall | 4 | 84.57 | 34.23 | 0.001 |
| Nest occupancy | Peak-season temperature + rainfall | 4 | 88.18 | 37.84 | 0.001 |
| Nest occupancy | Null | 2 | 50.34 | 0.00 | 0.993 |
| Nesting success | Pre-breeding MEI | 3 | 67.46 | 8.24 | 0.016 |
| Nesting success | Early-season temperature + rainfall | 4 | 84.98 | 25.77 | 0.001 |
| Nesting success | Entry temperature + rainfall | 4 | 85.61 | 26.40 | 0.001 |
| Nesting success | Peak-season temperature + rainfall | 4 | 94.19 | 34.97 | 0.001 |
| Nesting success | Null | 2 | 59.21 | 0.00 | 0.984 |

**Table S1. AICc model selection for nest occupancy and nesting success using pre-breeding MEI and local climate-window predictors.** Models were compared separately for occupancy and nesting success. *K* is the number of estimated parameters; ΔAICc is relative to the best-supported model for the corresponding response.

| Species | Year | Available nest-entry events | Missing temperature proportion | Retained for analysis |
|---|---|---|---|---|
| Great Hornbill | 2012 | 3 | 0.695 | No |
| Great Hornbill | 2013 | 5 | 0.351 | No |
| Great Hornbill | 2014 | 3 | 0.000 | Yes |
| Great Hornbill | 2015 | 6 | 0.000 | Yes |
| Great Hornbill | 2016 | 6 | 0.000 | Yes |
| Great Hornbill | 2017 | 6 | 0.000 | Yes |
| Great Hornbill | 2018 | 4 | 0.020 | Yes |
| Great Hornbill | 2019 | 3 | 0.874 | No |
| Great Hornbill | 2020 | 8 | 0.000 | Yes |
| Great Hornbill | 2021 | 7 | 1.000 | No |
| Great Hornbill | 2022 | 9 | 0.000 | Yes |
| Great Hornbill | 2023 | 5 | 1.000 | No |
| Great Hornbill | 2024 | 10 | 0.311 | No |
| Great Hornbill | 2025 | 9 | 0.450 | No |
| Wreathed Hornbill | 2012 | 4 | 0.695 | No |
| Wreathed Hornbill | 2013 | 6 | 0.351 | No |
| Wreathed Hornbill | 2014 | 7 | 0.000 | Yes |
| Wreathed Hornbill | 2015 | 6 | 0.000 | Yes |

| | | | | |
|---|---|---|---|---|
| Wreathed Hornbill | 2016 | 7 | 0.000 | Yes |
| Wreathed Hornbill | 2017 | 5 | 0.000 | Yes |
| Wreathed Hornbill | 2018 | 4 | 0.020 | Yes |
| Wreathed Hornbill | 2019 | 4 | 0.874 | No |
| Wreathed Hornbill | 2020 | 3 | 0.000 | Yes |
| Wreathed Hornbill | 2021 | 5 | 1.000 | No |
| Wreathed Hornbill | 2022 | 4 | 0.000 | Yes |
| Wreathed Hornbill | 2023 | 4 | 1.000 | No |
| Wreathed Hornbill | 2024 | 2 | 0.311 | No |
| Wreathed Hornbill | 2025 | 5 | 0.450 | No |
| Oriental Pied Hornbill | 2012 | 2 | 0.901 | No |
| Oriental Pied Hornbill | 2013 | 4 | 0.556 | No |
| Oriental Pied Hornbill | 2014 | 3 | 0.086 | Yes |
| Oriental Pied Hornbill | 2015 | 3 | 0.000 | Yes |

| | | | | |
|---|---|---|---|---|
| Oriental Pied Hornbill | 2016 | 3 | 0.000 | Yes |
| Oriental Pied Hornbill | 2017 | 3 | 0.000 | Yes |
| Oriental Pied Hornbill | 2018 | 3 | 0.020 | Yes |
| Oriental Pied Hornbill | 2019 | 4 | 0.682 | No |
| Oriental Pied Hornbill | 2021 | 11 | 1.000 | No |
| Oriental Pied Hornbill | 2022 | 2 | 0.000 | Yes |
| Oriental Pied Hornbill | 2023 | 3 | 0.801 | No |
| Oriental Pied Hornbill | 2024 | 1 | 0.106 | No |
| Oriental Pied Hornbill | 2025 | 3 | 0.245 | No |

**Table S2. Species-year audit of climate-data completeness and nest-entry records used in the sliding-window analysis.** "Missing temperature proportion" is the fraction of the expected daily temperature observations that were unavailable within the 150-day period preceding the species-specific reference date. It is calculated as the number of missing daily temperature values divided by the total number of calendar days in that 150-day window; values range from 0 to 1 and can be multiplied by 100 to express percentages. For example, 0.020 represents 2.0% missing daily temperature observations, whereas 0.695 represents 69.5% missing observations. A species-year was retained when the missing climate proportion was less than 0.10, equivalent to less than 10% missing daily observations. The table reports the number of nest-entry events available in each species-year and whether those events were retained for analysis. The retained dataset included 95 of 195 available nest-entry events from seven breeding seasons: 2014-2018, 2020, and 2022.

| Species | Model | *K* | AICc | ΔAICc | AICc weight | Species-years |
|---|---|---|---|---|---|---|
| Great Hornbill | Entry-window climate | 4 | 84.0 | 39.7 | <0.001 | 6 |
| Great Hornbill | Early-window climate | 4 | 83.8 | 39.5 | <0.001 | 6 |
| Great Hornbill | Peak-window climate | 4 | 78.8 | 34.5 | <0.001 | 6 |
| Great Hornbill | Null | 2 | 44.3 | 0.0 | 1.000 | 6 |
| Wreathed Hornbill | Entry-window climate | 4 | Inf | Inf | 0 | 5 |
| Wreathed Hornbill | Early-window climate | 4 | Inf | Inf | 0 | 5 |
| Wreathed Hornbill | Peak-window climate | 4 | Inf | Inf | 0 | 5 |
| Wreathed Hornbill | Null | 2 | 32.8 | 0.0 | 1.000 | 5 |
| Oriental Pied Hornbill | Entry-window climate | 4 | 86.4 | 39.1 | <0.001 | 6 |
| Oriental Pied Hornbill | Early-window climate | 4 | 87.2 | 39.9 | <0.001 | 6 |
| Oriental Pied Hornbill | Peak-window climate | 4 | 84.4 | 37.1 | <0.001 | 6 |

| | | | | | | |
|---|---|---|---|---|---|---|
| Oriental Pied Hornbill | Null | 2 | 47.2 | 0.0 | 1.000 | 6 |

**Table S3. AICc model selection for species-specific breeding duration.** For each species, breeding duration was modeled using a null model and three climate-window models based on the entry, early-season, and peak-season periods. *K* is the number of estimated model parameters. AICc is the small-sample corrected Akaike information criterion; ΔAICc is the difference between a model's AICc and the lowest AICc for that species; and AICc weight is the relative support for the model within that species' candidate set. The analysis included 6 species-years for Great Hornbill, 5 for Wreathed Hornbill, and 6 for Oriental Pied Hornbill. "Inf" indicates that the corresponding model fit produced an infinite AICc and therefore received zero AICc weight. No climate-window model had $\Delta AICc < 2$; therefore, no climate-window parameter estimate is interpreted as a supported effect.
